# Anesthetic oxygen use and sex are critical factors in the FLASH sparing effect

**DOI:** 10.1101/2023.11.04.565626

**Authors:** Armin D. Tavakkoli, Megan A. Clark, Alireza Kheirollah, Austin M. Sloop, Haille E. Soderholm, Noah J. Daniel, Arthur F. Petusseau, Yina H. Huang, Charles R. Thomas, Lesley A. Jarvis, Rongxiao Zhang, Brian W. Pogue, David J. Gladstone, P. Jack Hoopes

## Abstract

**Introduction:** Ultra-high dose-rate (UHDR) radiation has been reported to spare normal tissue compared to conventional dose-rate (CDR) radiation. However, reproducibility of the FLASH effect remains challenging due to varying dose ranges, radiation beam structure, and in-vivo endpoints. A better understanding of these inconsistencies may shed light on the mechanism of FLASH sparing. Here, we evaluate whether sex and/or use of 100% oxygen as carrier gas during irradiation contribute to the variability of the FLASH effect.

**Methods:** C57BL/6 mice (24 male, 24 female) were anesthetized using isoflurane mixed with either room air or 100% oxygen. Subsequently, the mice received 27 Gy of either 9 MeV electron UHDR or CDR to a 1.6 cm^2^ diameter area of the right leg skin using the Mobetron linear accelerator. The primary post-radiation endpoint was time to full thickness skin ulceration. In a separate cohort of mice (4 male, 4 female) skin oxygenation was measured using PdG4 Oxyphor under identical anesthesia conditions.

**Results:** In the UHDR group, time to ulceration was significantly shorter in mice that received 100% oxygen compared to room air, and amongst them female mice ulcerated sooner compared to males. However, no significant difference was observed between male and female UHDR mice that received room air. Oxygen measurements showed significantly higher tissue oxygenation using 100% oxygen as the anesthesia carrier gas compared to room air, and female mice showed higher levels of tissue oxygenation compared to males under 100% oxygen.

**Conclusion:** The FLASH sparing effect is significantly reduced using oxygen during anesthesia compared to room air. The FLASH sparing was significantly lower in female mice compared to males. Both tissue oxygenation and sex are likely sources of variability in UHDR studies. These results suggest an oxygen-based mechanism for FLASH, as well as a key role for sex in the FLASH skin sparing effect.

## 1. Introduction

Ultra-high dose-rate (UHDR) radiation has been shown to spare normal tissue (FLASH effect) as compared to conventional dose-rate (CDR) radiation ^1–3^. However, replication of the FLASH effect across institutions remains challenging ^4–7^. Studies demonstrating a positive FLASH effect prescribe increasingly limited – and at times conflicting – beam, dose, animal models, and endpoints to reproduce the effect. On the other hand, studies that show no FLASH sparing (or those that show a detrimental FLASH effect) are often dismissed for not having the correct UHDR “conditions” ^8^. This observation suggests that there are likely factors at play that are being left unaccounted for. These variations, in part, stem from the lack of a solidified understanding of the factors in the FLASH mechanism and how it is modulated by different variables such as the underlying biology of the model.

In CDR radiation therapy, it is widely believed that the presence of oxygen and generation of free radicals is necessary for optimal killing of cancer cells. This is supported by the observation that well-oxygenated tumors are 2-3 times more sensitive to radiation compared to hypoxic tumors ^9^. This radio-sensitization effect by oxygen, the oxygen enhancement ratio (OER), is calculated as the ratio of dose required to achieve the same biologic effect under hypoxic conditions compared to normoxic conditions. Notably, the OER varies by type of radiation therapy. For instance, OER with proton and photon therapy (OER ∼ 3) is about twice that of neutron or heavy-ion radiation therapy (OER ∼ 1.5) ^10^. The concept of OER raises several interesting questions: How do changes in tissue oxygenation modulate the FLASH effect, and how does the magnitude of this change vary based on types and doses of radiation?

Despite the mechanism(s) of the FLASH sparing effect being an active topic of research, some of the most prominent hypotheses behind the mechanism of FLASH sparing continue to be oxygen based ^11^. In fact, isolated studies have shown that hyperoxygenation and hypoxic conditions reduce or eliminate the FLASH effect ^12^. Yet, most of the FLASH literature fails to report on or control for in-vivo experimental variables that could meaningfully alter tissue oxygenation. These include type, concentration, and duration of anesthetic use, use of anesthetic oxygen, and physiological parameters such as body temperature, respiratory rate, and sex ^13^. This makes variations in tissue oxygen a likely and poorly controlled source of variability in UHDR studies. To our knowledge, there are no studies that evaluate the FLASH sparing effect in dermal tissue under room air and 100% oxygen conditions and between sexes, with direct tissue oxygen measurements.

*In vivo* measurements of tissue oxygenation are extremely challenging. While electrodes can provide precise readings, the measurement technique is damaging to the tissue, consequently not reporting the oxygenation of the normal healthy tissue, and only provides a point sample measurement ^14^. Other methods like paramagnetic oxygen sensors (EPR oximetry) are invasive and require injection-site healing, and NMR relaxation methods can only report on relative changes in averaged O2 levels ^15^. Therefore, optical methods to measure the quenching by oxygen of fluorescence probes with a sensor, either camera or fiber-optic fiber as the detector, have been used reliably to measure both distribution and average of the distribution, respectively, *in vivo*. The fluorescent probe PdG4 Oxyphor, specifically, has been used repeatedly to report extracellular oxygen levels in tissues, with the phosphorescence detector calibrated to extract absolute pO2 readings from the lifetime-based quenching of PdG4^16–18^.

Here, we hypothesized that anesthetic oxygen use and/or sex contribute to the variability of the FLASH effect, and aimed to evaluate this hypothesis using the mouse skin. We used subcutaneous injections of PdG4 Oxyphor to measure changes in tissue oxygenation under room air and 100% oxygen.

## 2. Materials & Methods

### 2.1 Skin Irradiation

#### Animals

Forty-eight C57BL/6 mice (24 male, 24 female), ranging from 8-10 weeks in age, were acquired from Jackson Laboratories and allowed to acclimate for at least 3 days. Two days prior to planned radiation delivery, mice were tagged, and the right leg and leg area was shaved.

On the day of radiation delivery, mice were anesthetized using isoflurane (induction: 3% isoflurane delivered at 500 ml/min for 3 minutes; maintenance: 1.5% isoflurane delivered at 100 ml/min) in either room air or 100% oxygen. Core body temperature was measured via a rectal probe and maintained at 37.5 degrees. Mice were, on average, maintained on anesthesia for 10 minutes prior to irradiation.

#### Radiation Delivery

A Mobetron intraoperative linear electron accelerator (IntraOp Inc. USA) was used to deliver a 27 Gy dose of 9 MeV CDR or UHDR radiation. A shielded collimator was used to collimate the radiation field to a 1.6 cm^2^ diameter circular field. The right leg of prone mice was positioned on a 3D printed holder and a 1.6 cm^2^ diameter area, centered at the mid-leg, was irradiated. For CDR delivery, the average dose rate was 0.17 Gy/s. For UHDR delivery, the average dose rate was 200 Gy/s, and the 27 Gy radiation was delivered in 2 pulses (2 × 3.16 us @120 Hz). Dosimetry was monitored by beam-current transformer measurements and verified by radiochromic film, calibrated from prior data.

#### Study Endpoint

Mice were checked daily for skin lesions at the irradiation site by two independently trained investigators. The day a full thickness ulcer was noted, lesions were photographed. Discrepancies in time to ulceration between the trained staff were resolved by a veterinary pathologist. Time to full thickness skin ulceration was used as the primary study endpoint for survival analysis. Mice that did not show lesions were monitored for 20 days post-irradiation and censored at that time for survival analysis. All non-lesion mice were monitored for an additional 10 days beyond the study endpoint to ensure lack of lesion development. The study design is summarized in figure 1.

**Figure 1.**
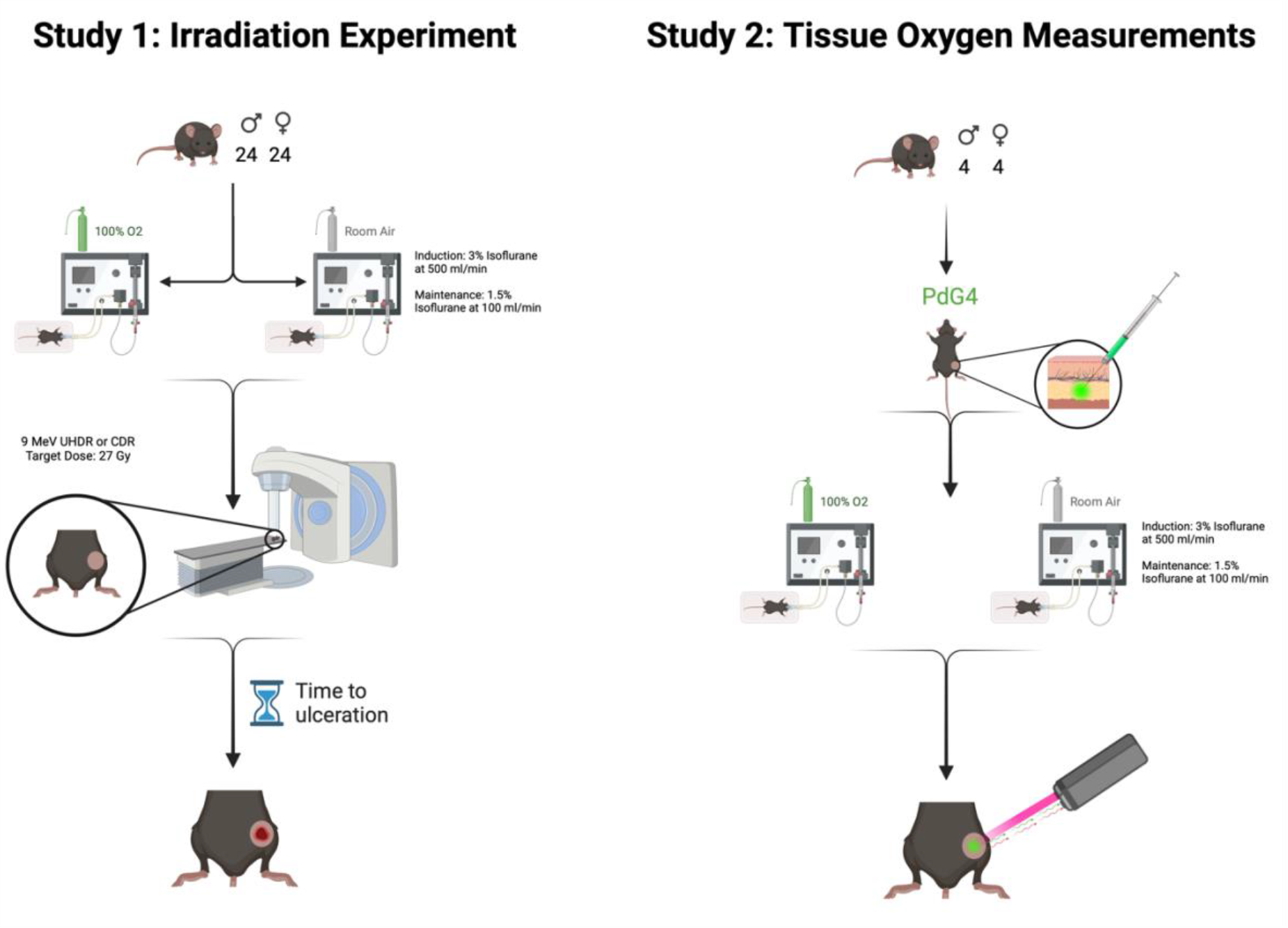
A visual demonstration of the study design. In study 1, forty-eight C57BL/6 mice (24 male, 24 female) were anesthetized using isoflurane delivered in 100% oxygen or room air and received 27 Gy of radiation to right leg skin using a Mobetron linear accelerator. Time to skin ulceration was measured as the primary endpoint. In study 2, eight C57BL/6 mice (4 male, 4 male) received subcutaneous injections of PdG4 Oxyphor and were anesthetized similar to study 1. An excitor-detector fiber-optic pair was used to read tissue oxygenation levels at the injection site.

### 2.2 Measuring Tissue Oxygenation

Eight C57BL/6 mice (4 males, 4 females) were used over the course of two days; on the first day, the mice were anesthetized with 100% oxygen mixed with isoflurane and on the second day the mice were induced while breathing room air mixed with isoflurane. To simulate the breathing conditions of the mouse on the day of irradiation, each mouse was anesthetized as described in section 2.1. After a 3-minute induction period, the left rear leg of the mice was shaved and 0.05mL of 100uM PdG4 dissolved in phosphate-buffered saline (PBS) was injected subcutaneously into the leg of the mice. The phosphorescence lifetime was read out by a commercial system (OxyLED, Oxygen Enterprises, Philadelphia PA) that was calibrated with the Stern-Volmer emission time constants for the PdG4 sample injected. From this system, the fiber pair contained a pulsed red (637nm) excitation light and a collecting fiber, connected to an APD detector. This fiber was positioned approximately 5mm from the skin surface and oxygen pressure (mmHg) was readout repeatedly for ten minutes to sample the tissue pO2 value. Core body temperature was maintained at 37.5 degrees via external heating pad.

### 2.3 Statistical Analyses

Using time to skin ulceration as the primary study endpoint, Kaplan-Meier survival curves were constructed for the irradiated mice. The log-rank (Mantel-Cox) were used to compare survival data and verified by Cox regression tests. The tissue oxygen measurement data was analyzed using independent sample t-tests. All analyses were conducted using the SPSS Statistical Package (IBM Corp., USA). Graphics were constructed using GraphPad Prism (Prism Inc, USA) and BioRender software.

## 3. Results

### 3.1 Skin Irradiation

Actual radiation dose delivered at the surface was 27 Gy ± 1.5 Gy (5%). All mice were checked daily for skin lesion development at irradiation site. As we have previously described, mice typically progress through dry and wet skin desquamation prior to full thickness ulceration/epidermolysis, though detection of wet and dry squamation can be variable. Therefore, time to full thickness skin ulceration was used at the primary endpoint.

Within the UHDR group, out of 12 mice breathing room air, 5 developed full thickness ulceration. The remaining 7 mice did not develop lesions during the 20-day study period, nor the 10 day follow up period, and were therefore censored from survival analyses at 20 days. Therefore, the median survival (time to skin ulceration) for room air breathing UHDR mice is greater than 20 days. On the other hand, out of the 12 mice breathing 100% oxygen, 11 developed ulceration and 1 did not, with a median time to skin ulceration of 12 days. This difference in time to skin ulceration was significantly different between the groups, p<0.05.

Within the CDR group, out of 12 mice breathing room air, 11 developed ulcers with a median time to ulceration of 9.5 days. Likewise, out of the 12 mice breathing 100% oxygen, 11 developed ulcers, but with a median time to ulceration of 15.5 days. This difference was not significantly significant.

Comparing the UHDR to CDR group, UHDR mice breathing 100% oxygen did not significantly differ from CDR mice breathing 100% oxygen in median time to skin ulceration of 12 and 15.5 days, respectively. Conversely, UHDR mice breathing room air showed significantly longer time to ulceration compared to CDR mice breathing room air, with median time to skin ulceration being greater than 20 days compared to 9.5 days, respectively. This data is summarized in Figure 2.

**Figure 2.**
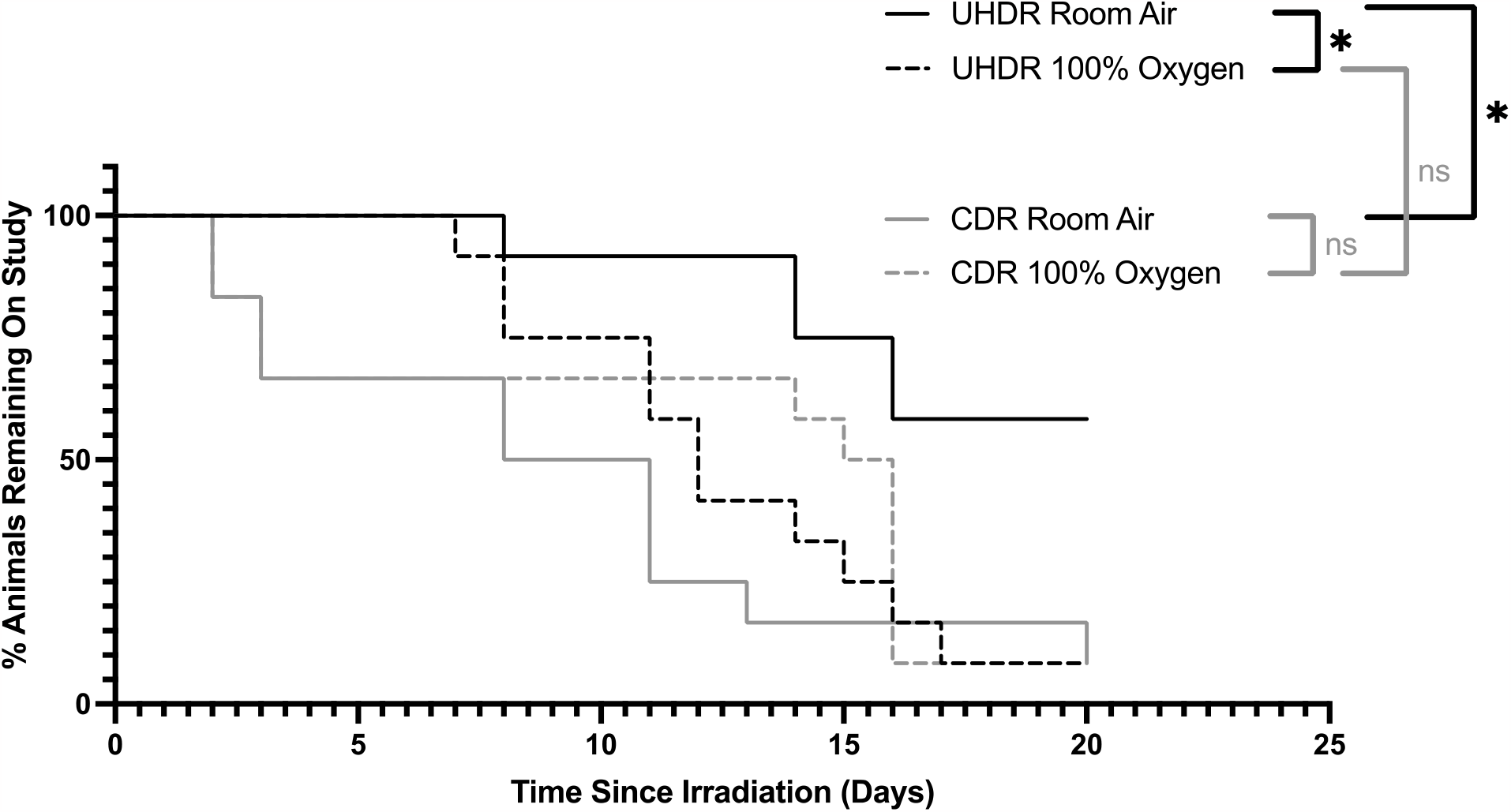
Kaplan-Meier curves demonstrating time to skin ulceration in leg skin of UHDR and CDR irradiated mice under room air and 100% oxygen conditions. Animals were removed from the study when a full thickness ulcer developed. Mice that did not show lesions after 20 days post-irradiation were censored for survival analysis and monitored for an additional 10 days to ensure no lesions developed. UHDR room air mice (median = >20 days) developed ulcers significantly later than both UHDR 100% oxygen (median = 12 days) and CDR room air mice (median = 15.5 days). UHDR 100% oxygen mice did not differ significantly from CDR 100% oxygen mice (median = 12 days). CDR room air and CDR 100% oxygen mice did not differ from each other in time to ulceration. * = p < 0.05, ns = not statistically significant.

Separating the irradiation data by sex provided further insight into these differences. Within the UHDR group, female and male mice breathing room air did not differ significantly in time to skin ulceration, with median time to ulceration of greater than 20 days compared to 18 days, respectively. In contrast, female mice breathing 100% oxygen developed skin ulceration in a median of 11 days, which was significantly shorter than male mice breathing 100% oxygen with a median time to ulceration of 15.5 days. This is summarized in Figure 3A.

**Figure 3.**
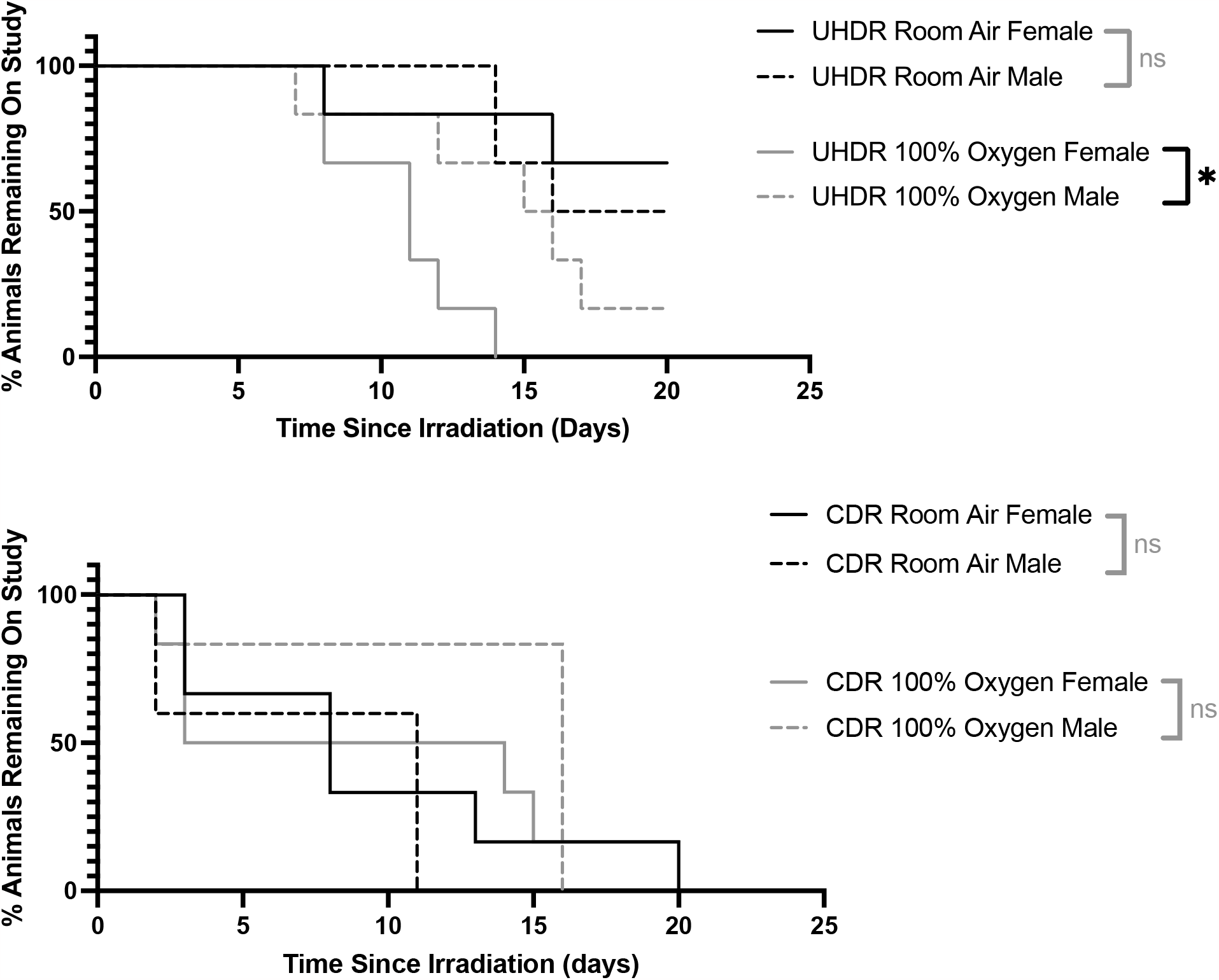
Kaplan-Meier curves demonstrating sex differences in time to skin ulceration in leg skin of UHDR and CDR irradiated mice under room air and 100% oxygen conditions. (**A**) Male and female UHDR room air mice did not differ in time to skin ulceration. However, female mice in UHDR 100% oxygen developed ulcers earlier compared to male mice in the same group. (**B**) No significant sex differences were seen between male and female CDR mice in neither room air nor 100% oxygen conditions. * = p < 0.05, ns = not statistically significant

Looking at sex differences in the CDR group, female and male mice breathing either room air or 100% oxygen did not differ significantly in time to skin ulceration. This data is summarized in Figure 3B.

### 3.2 Skin Oxygenation Measurements

Mean pO2 of the male mouse leg while breathing 100% oxygen (36±7mmHg) was significantly higher than that for the males breathing room air (21±3mmHg). Similarly, the mean pO2 of the female mouse leg while breathing 100% oxygen (56±11mmHg) was significantly higher than that for the females breathing room air (26±4mmHg). Interestingly, the mean pO2 for males as compared to females breathing 100% oxygen were significantly different from each other, as well; however, the difference when breathing room air was not significant. Results of tissue oxygenation measurements are summarized in Figure 4.

**Figure 4.**
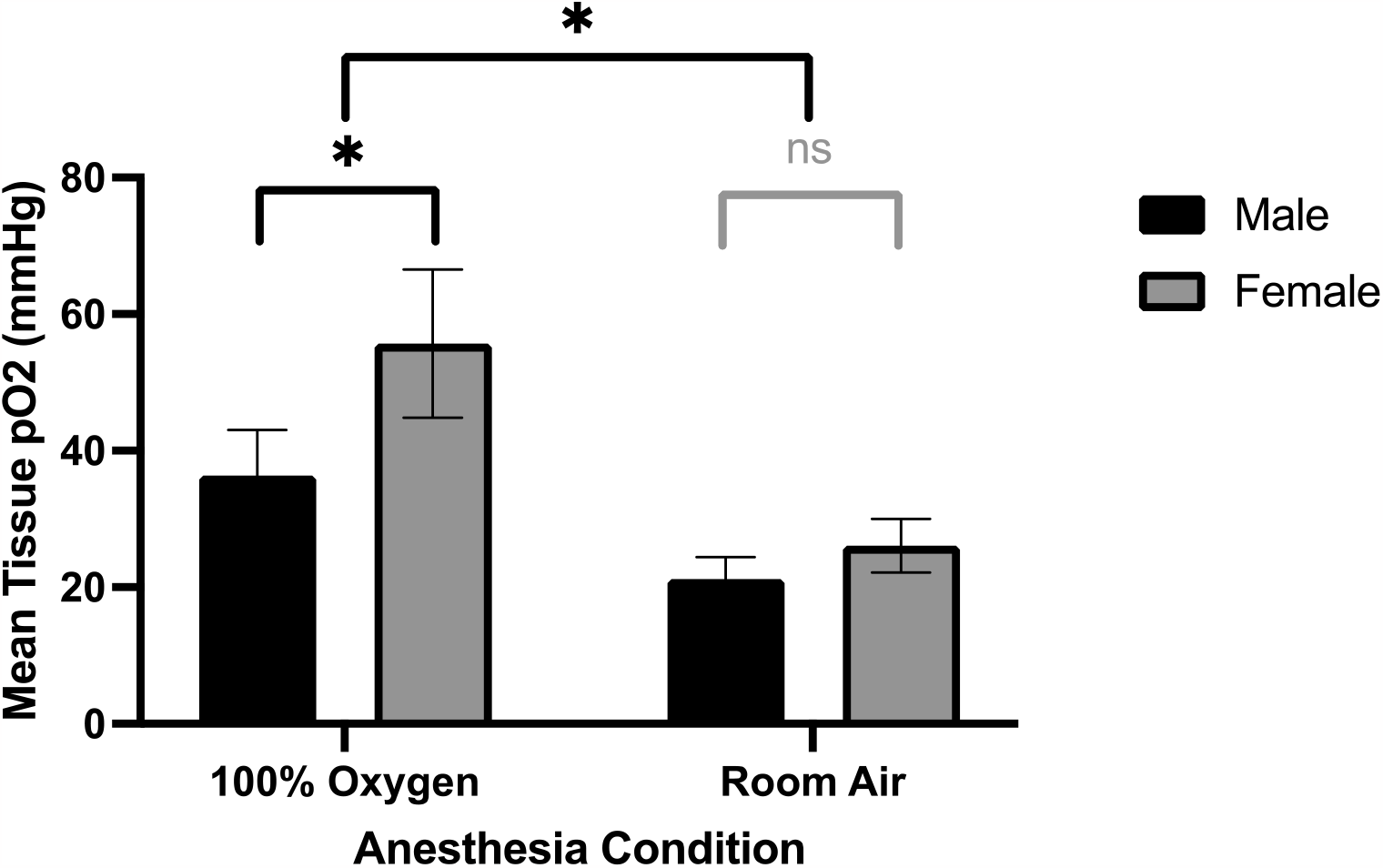
Direct tissue pO2 measurements under room air and 100% oxygen conditions using PdG4 Oxyphor. Mice that received 100% oxygen had significantly higher tissue pO2 levels compared to room air mice. The female mice that received 100% oxygen (mean = 56 mmHg, SD = 11) had higher tissue pO2 levels compared to males that received 100% oxygen (mean = 36 mmHg, SD = 7). * = p < 0.05, ns = not statistically significant.

## 4. Discussion

The literature on UHDR radiation and the FLASH effect shows wide variability in presence and extent of normal tissue sparing, hinting that there may be unknown sources of variability in in-vivo studies. Here, we set out to determine the effect of oxygen use during anesthesia on FLASH sparing of normal skin tissue, supplemented by tissue oxygen measurements. We accomplish this by irradiating animals with CDR or UHDR radiation while delivering anesthesia using either 100% oxygen or room air as carrier gas and measuring time to skin ulceration.

Our results show that time to ulceration in CDR radiation is not significantly affected by oxygen concentration of carrier gas. Conversely, UHDR irradiated mice that received room air showed a significantly increased time to skin ulceration. Indeed, over half of the mice in the UHDR room air group did not show ulceration over the course of the study, compared to just one mouse in the UHDR 100% oxygen group. Comparing the UHDR to CDR irradiated mice, we show no FLASH sparing effect in mice breathing 100% oxygen. On the other hand, significant sparing was seen between UHDR and CDR mice receiving room air as carrier gas.

Breaking down our results by type of radiation and sex, we show a significantly reduced time to ulceration in female mice that received UHDR while breathing 100% oxygen compared to male. However, no sex difference was seen between male and female mice that received UHDR radiation while breathing room air, nor CDR mice under either condition. This is an interesting finding, particularly considering sex differences seen in our tissue oxygen measurements.

Using the PdG4 Oxyphor, we were able to repeatably measure oxygen levels in the mouse leg skin tissue. Although tissue oxygenation levels are similar between male and female mice at room air, female mice breathing 100% oxygen show significantly higher tissue pO2 levels compared to male mice breathing 100% oxygen. This difference correlates with that seen in time to ulceration between male and female mice in UHDR 100% oxygen group, where females (who showed higher tissue oxygen levels) ulcerate faster and are thus more radiosensitive. Though the mechanism behind higher tissue oxygenation in females requires further investigation, we propose higher estrogen levels in female mice as a possible contributor. Estrogen is a potent angiogenesis factor, and it plays a critical in both maintenance and repair of dermal blood vessels ^19^. If the dermis of the female mouse effectively vascularized, higher tissue oxygenation under anesthesia would be expected.

In summary, we make two important observations: first, presence of 100% oxygen negated any FLASH sparing effect, primarily through its effect on UHDR irradiated mice; oxygen did not affect time to ulceration in the CDR irradiated mice. Second, within the UHDR irradiated mice, male and female mice were equivalent in their time to ulceration under room air conditions but differed significantly from each other under 100% oxygen condition. This is in direct agreement with the tissue oxygen measurements previously presented. However, once again, this difference was not seen in CDR irradiated mice.

We suspect other anesthetic conditions, such as depth of anesthesia (isoflurane/sevoflurane concentration, flow rate, length of induction and maintenance) and physiologic parameters such as respiratory rate and body temperature are likely to have a pronounced effect on tissue oxygen levels, and consequently the extent of sparing in UHDR radiation. For instance, in a current, separate study, we find that maintaining animals at 3% isoflurane results in significantly lower tissue oxygenation compared to the 1.5% used in this study.

Further investigation into the source of the sex differences in FLASH sparing and tissue oxygenation is also needed. Skin as a model for FLASH effects may also be problematic because of the variability in control over skin oxygenation between animals, sex, and physiologic conditions, although at the same time it presents as an interesting model in which factors affecting the mechanism of FLASH may be teased out. This work makes clear the need for careful control and reporting of these variables in future UHDR in-vivo studies.

## 5. Acknowledgements

We are grateful for the radiation resources provided by Dartmouth-Hitchcock Medical Center Department of Radiation Oncology.

